# Comprehensive profiling of wastewater viromes by genomic sequencing

**DOI:** 10.1101/2022.12.16.520800

**Authors:** Emanuel Wyler, Chris Lauber, Artür Manukyan, Aylina Deter, Claudia Quedenau, Luiz Gustavo Teixeira Alves, Stefan Seitz, Janine Altmüller, Markus Landthaler

## Abstract

Genomic material in wastewater provides a rich source of data for detection and surveillance of microbes. Used for decades to monitor poliovirus and other pathogens, the SARS-CoV-2 pandemic and the falling costs of high-throughput sequencing have substantially boosted the interest in and the usage of wastewater monitoring. We have longitudinally collected over 100 samples from a wastewater treatment plant in Berlin/Germany, from March 2021 to July 2022, in order to investigate three aspects. First, we conducted a full metagenomic analysis and exemplified the depth of the data by temporal tracking strains and to a certain extent also variants of human astroviruses and enteroviruses. Second, targeting respiratory pathogens, a broad enrichment panel enabled us to detect waves of RSV, influenza, or common cold coronaviruses in high agreement with clinical data. Third, by applying a profile Hidden Markov Model-based search for novel viruses, we identified more than 100 thousand novel transcript assemblies likely not belonging to known virus species, thus substantially expanding our knowledge of virus diversity. Taken together, we present a longitudinal and deep investigation of the viral genomic information in wastewater that underlines the value of sewage surveillance for both public health purposes and planetary virome research.

## Introduction

Environmental surveillance of wastewater samples combined with high-throughput genomic sequencing can be a particularly valuable tool to monitor the vast diversity of microbes and anti-microbial resistance genes (Leifels et al., 2022; Santiago-Rodriguez, 2022). Accumulating genomic data from environmental samples can serve three purposes. First, it can be used as a sentinel system to monitor human pathogens and inform public health decisions (Diamond et al., 2022). Wastewater was and is essential for assessing Polio outbreaks (Anis et al., 2013; Chowdhary and Dhole, 2008; Ryerson et al., 2022). However, a range of other viruses can be detected, including gastroenteritis and hepatitis viruses, and influenza (Heijnen and Medema, 2011; Hellmer et al., 2014). In the SARS-CoV-2 pandemic, wastewater monitoring was established worldwide to monitor the virus in almost real time (Diamond et al., 2022).

The second use is to characterize microbial community ecosystems, which can be indicators for e.g. quality and function of freshwater systems (Numberger et al., 2022). And third, the advent of ultra-deep sequencing and computational methods, so far from public databases and seawater samples, has enabled the detection of large number of novel viruses, thus expanding considerably our knowledge of viral diversity (Edgar et al., 2022; Gregory et al., 2019; Martinez-Hernandez et al., 2022).

For monitoring human viruses in wastewater, various aspects need to be considered. This includes how much and in which form genomic material is being shed. Whereas in stool samples, for example, substantial amounts of infectious enteroviruses are found (Blomqvist and Roivainen, 2016), it is debated whether e.g. SARS-CoV-2 RNA in feces is present as part of intact viral particles or – what may be more prevalent – as fragments (Cerrada-Romero et al., 2022; Guo et al., 2021b). Facing the general higher stability of non-enveloped viruses (picornaviridae such as enteroviruses, caliciviruses or astroviruses) when compared to enveloped viruses (influenza, coronaviruses, RSV etc.) (Firquet et al., 2015), this can lead to substantial differences in the detection potential in addition to the incidence rates in the population. Overall, human pathogens are only a minor part of the totality of microbes in wastewater (Cantalupo et al., 2011; Wu et al., 2019), making their detection challenging. It is also important to note that distinct viral nucleic acids can be present in different fractions of wastewater samples. For example, both mpox and influenza genomic nucleic acids have been found to be associated with solids (Mercier et al., 2022; Wolfe et al., 2022b).

Previous studies have shown that wastewater is a useful source to detect circulating known as well as novel viruses (Adriaenssens et al., 2018; Bibby and Peccia, 2013; Cantalupo et al., 2011; Fernandez-Cassi et al., 2018; Guajardo-Leiva et al., 2020; Martinez-Puchol et al., 2021; Perez-Cataluna et al., 2021; Rothman et al., 2021). These studies explored several aspects, including enrichment of specific viruses (Martinez-Puchol et al., 2021; Rothman et al., 2021). Whereas most studies analyzed a small number of samples, a recent investigation collected 85 samples over a period of five months (Rothman et al., 2021), showcasing the relevance of longitudinal samplings.

In our work – although we all superkingdoms as well as antimicrobial resistance genes in the data, we focused our analysis on viruses –, we extend this body of work in several directions. First, we covered a time period of one and a half years with 116 samples in total, including a time period with reduced hygiene measurements compared to the height of the SARS-CoV-2 pandemic. The longitudinal sampling allows the observation of seasonal recurrence and monitoring of clinically relevant viruses such as respiratory syncytial virus (RSV). Second, very deep sequencing reveals subspecies of several viruses (exemplified with mamastroviruses and enteroviruses), and, for astroviruses, also enables to track virus variant development over the sampling time period by assessing single point mutations. Third, the enrichment method applied here shows high concordance between clinical sampling and wastewater data for several respiratory viruses. And fourth, we substantially extend knowledge about viral diversity by identifying more than 100 thousand contigs belonging to previously unknown RNA and DNA viruses. In summary, our study shows that wastewater is an extraordinary rich source of nucleic acids to track known and novel viruses.

## Methods

### Sample collection

The sample collection was done as described previously (Schumann et al., 2022), or as following. Samples were collected (except for the first date, see supplementary table S1) from a single wastewater treatment plant in Berlin, Germany, as two hours composite samples (8–10 pm and 10–12 pm) at the primary influent collector, except for the indicated 24 hours composite samples on July 23, 2022. Berlin wastewater treatment plant effluents usually contain 500–1500 mg/L chemical oxygen demand, 200–600 mg/L suspended solids, 40–80 mg/L ammonium-N, 2–8 mg/L orthophosphate-P, 1500–2000 μS/cm electrical conductivity.

### Sample processing and RNA isolation

The sample processing was essentially done as described previously (Schumann et al., 2022), with modifications for some samples. Samples were kept at four degrees, until processed about 12 h after collection. Processing was done along a published protocol (Jahn et al., 2022). First, the raw sample was filtered through 2 µm glass fiber and 0.2 μM PVDF filters (Millipore, cat# AP2007500 and S2GVU02RE). For the standard procedure, 60 ml filtrate were subsequently concentrated on a 10 kDa cutoff centricon unit (Millipore, cat# UFC701008), that was previously pre-conditioned with 50 mL ultrapure water centrifuged with 3000 g for 15 minutes at 4 °C, for 30 minutes at 3000 g/4 °C. The concentrate (about 300-450 µl) was mixed 1:3 with Trizol LS (ThermoFisher cat# 10296-010), and the RNA extracted using the DirectZol RNA kit (Zymo cat# R2052), including DNase treatment, and eluted in 50 μL ultrapure water according to the manufacturer’s instruction. When using nanotrap beads (Ceres Nanosciences, cat# 44202), 10 ml filtered wastewater were mixed with 100 µl Enhancement Reagent 2, followed by addition of 150 µl beads, following extraction according to the manufacturer’s protocol. RNA was subsequently extracted using either Trizol/DirectZol or the ZymoBIOMICS RNA/DNA combination extraction (Zymo, cat# ZYM-R2002).

### Reverse transcription and quantitative PCRs

For the RT-qPCRs, 16 µl RNA were mixed with 4 µl LunaScript master mix (NEB cat# E3010L), according to the manufacturer’s instructions, except for a 20 min incubation at 55 °C instead of 10 min. Afterwards, the cDNA was diluted 1:10 with ultrapure water, and 3.75 μL diluted cDNA used per qPCR reaction, using a SYBR green master mix (ThermoFisher cat# 43-643-46), with 250 nM final concentration of the primers listed in Supplementary Table S2.

### Enrichment using ElementZero beads

SARS-CoV-2 and TBRV RNA, respectively, were enriched from total RNA using the SARS-CoV-2 / TBRV MagIC beads (ElementZero Biolabs) according to the manufacturer’s protocol.

### Total RNA sequencing

RNA sequencing were prepared using the SMARTer® Stranded Total RNA-Seq Kit v3 -Pico Input Mammalian (Takara cat# 634485) with 5 µl RNA from either the total RNA or the ElementZero eluate as starting material according to the manufacturer’s protocoll. Libraries were then pooled and sequenced on a Novaseq 6000 device with 2×109 bp paired-end sequencing.

### Enrichment using the xHYB adventitious agent panel

Sequencing libraries were pooled according to Supplementary Table S1 into three pools, with an approximate equal amount of starting material for every sample. The pools were then processed individually using the Qiagen xHYB adventitious agent panel (Qiagen cat# 333355) according to the manufacturer’s protocol, and sequenced after the final re-ampflication step.

### Metagenomic analysis using kaiju

Total RNA-seq data was analyzed by the kaiju program (Menzel et al., 2016). For both the application of kaiju pipeline as well as the subspecies analysis, the code will be detailed in the github repository. Specifically, the cd-hit-dup command from CD-HIT (Fu et al., 2012) was used to filter duplicated reads out, and kaiju command was incorporated to assign taxonomies to remaining reads from each 116 samples. Custom R scripts were used to summarize duplicated and assigned reads. Initial low count filtering was conducted by removing annotations with a maximum count of 10 across all samples. We have used Hellinger transformation (Nieuwenhuijse et al., 2020) to normalize the count table. Principal components analysis was performed on the normalized counts, and top PC1 and PC2 loadings were used to detect annotations that are most correlated with PC1 and PC2. Additional low count filtering removed annotations with mean count lower than 10 which revealed an additional set of annotations that are associated with outlying samples (i.e. outlying annotations). The following R packages were used for data processing and visualization: dplyr (Wickham, 2022b), ggplot2 (Wickham, 2022a), Complex heatmaps (Gu et al., 2016), pheatmap (Kolde, 2019), msa (Bodenhofer et al., 2015), reshape2.

### Analysis of viral abundances and variants

Alignements were done using hisat2 (Kim et al., 2019), and read counting performed using samtools (Danecek et al., 2021).

### Sequence contig assembly

Adapter sequences and low-quality bases were trimmed from the raw sequencing reads using fastp v0.23.2 (Danecek et al., 2021) with parameters ‘-q 20 --dedup’. The trimmed reads were assembled into scaffolds in paired-end mode using SPAdes v3.15.4 (Prjibelski et al., 2020) with default parameters. Each of the 116 RNA sequencing experiments was assembled separately.

### Taxonomic classification of sequences

We extracted peptide sequences encoded by open reading frames (ORFs) of at least 300 nucleotides in length from the scaffolds using getorf from the EMBOSS package v6.6.0.0 (Rice et al., 2000). The peptide sequences were classified at the taxonomic ranks superkingdom, kingdom, phylum, subphylum, class, order, suborder, family, subfamily, genus, subgenus and species using the MMseqs2 taxonomy module v5f8735872e189991a743f7ed03e7c9d1f7a78855 (Hauser et al., 2016). We used the full nr database, downloaded in March 2022, for this analysis. Sequences classified as Bacteria, Eukaryota or Archaea at the superkingdom rank and the scaffold sequences they originated from were not considered further.

### Discovery of viral sequences

We applied the following multi-stage process to identify and annotate viral sequences in the set of assembled scaffolds. We run a profile Hidden Markov Model (pHMM)-based sequence homology search against predicted peptide sequences encoded by ORFs of at least 300 nucleotides in length using hmmsearch from the HMMER v3.1b1 package (Eddy, 2011) in default mode. We used the following sets of pHMMs: the combined set of 84420 profiles from VirSorter 2 (Guo et al., 2021a), a set of 74 lineage major capsid protein (MCP) profiles of nucleocytoplasmic large DNA viruses (NCLDVs) (Schulz et al., 2020), five RNA-dependent RNA polymerase (RdRp) profiles of putative novel RNA virus phyla from the Tara Oceans Virome project (Zayed et al., 2022), 8390 profiles from the RNA Virus in MetaTranscriptomes (RVMT) project (Neri et al., 2022), 68 RdRp profiles from RdRp Scan v0.90 (Charon et al., 2022), and several in-house pHMMs of DNA and RNA virus proteins.

From the hits obtained during the pHMM searches we kept those sequences that were not classified as Eukaryota, Bacteria or Archaea at the superkingdom level by MMseqs2. To remove sequence redundancy, we clustered the scaffolds at 95% nucleotide sequence identity using MMseqs2 easy-linclust with parameter ‘--min-seq-id 0.95 -c 0.65 --cluster-mode 2’. To assess the genetic distance of these non-redundant, putatively viral sequences we run a DIAMOND blastx v2.0.13.151 (Buchfink et al., 2021) search with parameters ‘--ultra-sensitive –masking 0 -k 1 -f 6’ against the following set of known viral sequences: 590872 viral proteins from NCBI RefSeq (O’Leary et al., 2016), 311725 RdRp sequences from the Serratus and PalmDB projects (Babaian and Edgar, 2021; Edgar et al., 2022), 49421 RdRp footprint sequences from the Tara Oceans Virome project (Zayed et al., 2022), 77510 RdRp sequences from RVMT (Neri et al., 2022) and 15081 RdRp sequences from RdRp Scan v0.90 (Charon et al., 2022).

### Estimation of viral abundance

The trimmed sequencing reads were aligned in paired-end mode to the viral scaffolds using Bowtie 2 v2.3.4.1 (Langmead and Salzberg, 2012) with parameters ‘--no-unal -L 20 -N 1’. Samtools v1.10 (Danecek et al., 2021) was used to sort and index the resulting SAM/BAM files and to count the number of aligned reads per scaffold. Virus abundance was calculated as the total number of reads aligning to a scaffold across all sequencing libraries divided by the scaffold length. In comparisons between sequencing libraries, abundance was calculated as the number of reads of a particular library aligning to the viral scaffold divided by scaffold length divided by the total number of reads in the library. Abundance of a certain virus taxon (for instance a virus family or order) was calculated as the sum of the abundance values of all scaffolds classified as belonging to this taxon.

### Phylogenetic analysis of novel *Bunyavirales* sequences

We selected all scaffolds classified as *Bunyavirales* or the sister order *Articulavirales*. To reconfirm correct classification of these sequences, we conducted a pHMM search against profiles that we constructed based on 24 order-level RdRp alignments obtained from the Serratus project (Edgar et al., 2022) as well as in-house glycoprotein (GP) and nucleocapsid protein (NP) profiles of all 13 recognized virus families of the order *Bunyavirales*. We only kept sequences that showed the lowest E-value against either *Bunyavirales* or *Articulavirales* profiles in this search.

Putative RdRp protein sequences were aligned using MAFFT v7.310 (Katoh and Standley, 2013) with parameters ‘--localpair --maxiterate 1000 --reorder’ followed by manual curation. For phylogenetic tree reconstruction, *Mononegavirales RdRp* sequences available at NCBI RefSeq, clustered at 20% amino acid sequence identity using MMseqs2, were added as an outgroup. We also added *Bunyavirales and Articulavirales* reference proteins, clustered at 90% amino acid sequence identity. In addition, we included bunya-und articulavirus sequences that we discovered in an independent screen of eukaryotic transcriptome projects in the Sequence Read Archive (SRA); details of the method are described here (Lauber et al., 2021) and a general introduction to this type of data-driven virus discovery approach is reviewed here (Lauber and Seitz, 2022). We only considered 16 out of more than 8000 potential bunya-or articulavirus-like contigs from our SRA search that covered a sufficient length of the RdRp and which showed at least 80% protein sequence identity to one of the bunya-or articulavirus sequences retrieved from the wastewater data. The fact that we re-discovered some of the viral sequences from the wastewater analysis in the SRA analysis provides independent support for the authenticity of the described novel viruses.

We reconstructed a Bayesian phylogenetic tree using BEAST v1.8.0 (Suchard et al., 2018) with the LG+G4+I substitution model, a relaxed molecular clock model with lognormally distributed rates and a Yule speciation tree prior. Two chains were run for 5 million generations and their convergence was verified using Tracer (Rambaut et al., 2018) after removing the first 10% of sampled trees as burn-in. The maximum clade credibility tree was visualized in R using ggtree v3.4.0 (Yu, 2020).

## Results

### Sample collection and processing

We aimed to perform a deep and longitudinal profiling of microbes and particularly viruses in wastewater. To this end, we collected raw influx samples between March 2021 and July 2022 from a wastewater treatment plant in Berlin/Germany, using a procedure that depletes intact bacteria by including a 0.2 µm filtration step (Fig. 1A). Of note, SARS-CoV-2 characterization in some of these samples has been described previously (Schumann et al., 2022). A full overview of the samples including Ct values from RT-qPCR assays on various viruses is shown in Supplementary Table S1. Following RNA isolation, we generated 116 high-throughput sequencing libraries, which yielded in total 1.45 billion 2×109 bp read pairs (12 million on average per sample), with about 10-30% of reads duplicated (Fig. S1A).

**Figure 1.**
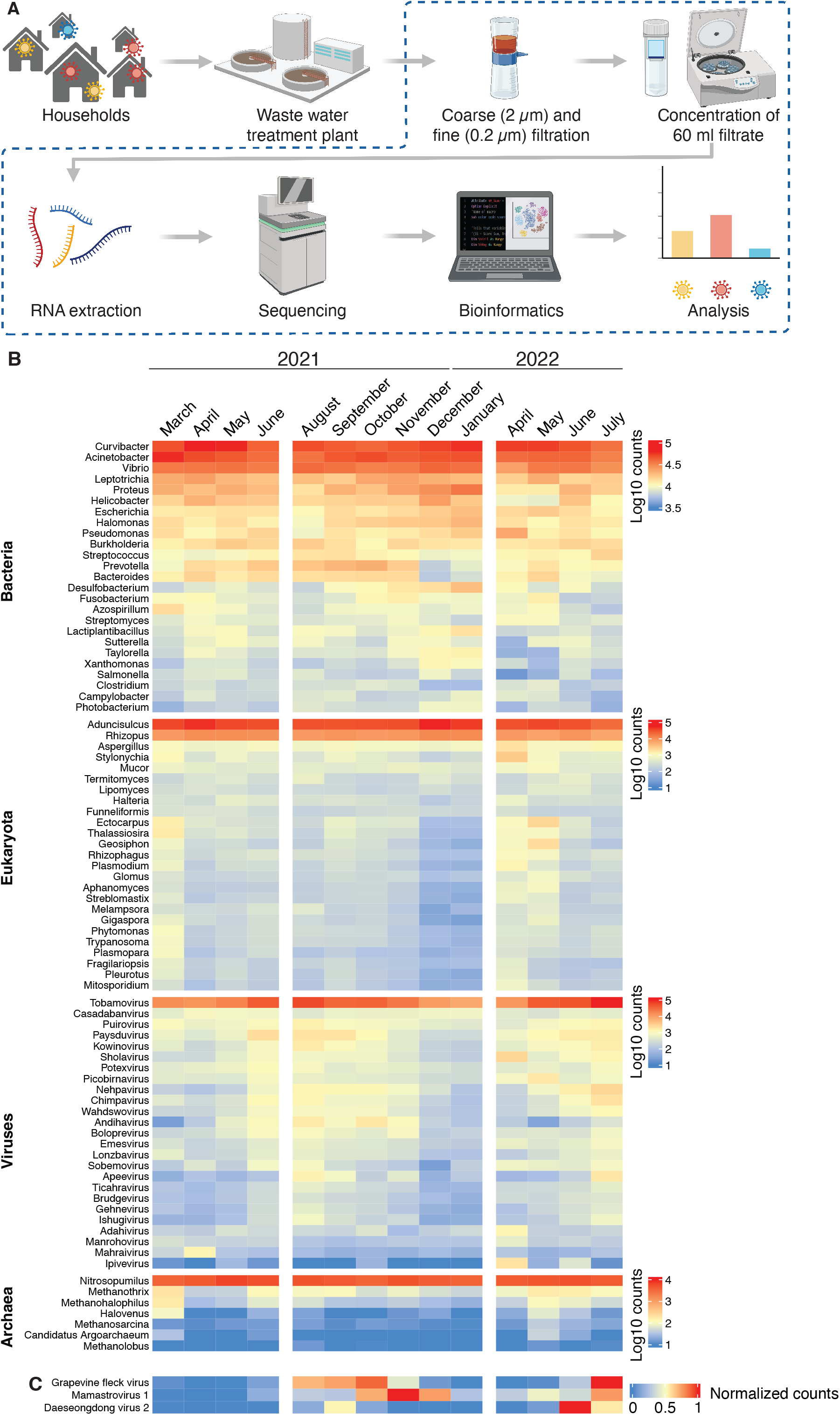
Diversity of microbes identified in wastewater and seasonality of viruses. **A**, overview of the sample processing pipeline. Figure created with BioRender.com. **B**, heatmap depicting abundance of the top genera for the three superkingdoms and viruses. Reads are shown as log10 transformed mapped per million, aggreated per month as indicated in the bottom. **C**, signal for three specific viruses, normalized on a 0-1 scale.

Most of the samples were processed using the standard procedure (Fig. 1A). In addition, some samples were processed using the Ceres nanotrap beads (see Supplementary Table S1 and method section for details). Except if indicated, for the following analysis only sequencing data from the samples with the standard procedure is used. Note that for some months, none or only one or two samples were successfully processed, henceforth they are omitted in some of the analysis shown below.

### A wide range of eukaryotes, bacteria, and viruses can be detected, however only viruses show specific patterns

For an initial assessment of the diversity of the detected organisms in the sample, we applied the metagenomics pipeline kaiju (Menzel et al., 2016). As in previous wastewater metagenomics studies (Rothman et al., 2020), bacteria were more abundant despite their depletion by size-exclusion (Fig. S1B). Initially, kaiju annotated reads to 67322 taxonomies. Low count filtering on all samples preserved 16082 taxonomies. A principal components analysis (Fig. S1C) using these taxonomies revealed an additional set of 11984 taxonomies (Methods) that are detected only in very few (1-3) samples. These highly variable taxonomies had particularly high read counts in the outlier samples (Fig. S1D).

Analyzing the 16082 taxonomies, we found genera from all three superkingdoms and viruses, with some however clearly dominating the dataset (Fig. 1, Fig. S1E). Of note, a recently published metagenomic study of wastewater treatment plant influxes also in Berlin/Germany from an earlier time period, has also found Acinetobacter among the most abundant bacterial species (Numberger et al., 2022).

Furthermore, when comparing our dataset with a recently defined “baseline for global urban virome surveillance in sewage” (Nieuwenhuijse et al., 2020), mostly the same virus families were the most abundant ones, such as *Virgaviridae, Siphoviridae, Astroviridae, Myoviridae, Dicistroviridae, Podoviridae, Microviridae, and Picornaviridae* (Fig. S1F).

In order to investigate individual virus species, we quantified relative levels of viruses from seasonal food, a human pathogen previously identified to be abundant in wastewater, and a virus infecting the common house mosquito, present in Berlin only during the Northern hemisphere summer. For the grapevine fleck virus, and for the Daeseondong virus 2 (host C. pipiens), we saw the expected patterns with peaks only in late summer/fall. Mamastrovirus 1, a virus causing gastroenteritis in humans (Boujon et al., 2017), also showed a distinct pattern. Overall, most of the sequencing reads belonged to a limited number of genera without apparent temporal patterns that recapitulates previous waste water metagenomic studies. Single virus species however showed specific patterns, which we also took as an indication that the overall sampling and data analysis was appropriate.

### Strain distribution and variant emergence of astroviruses

As observed in the data, astroviruses were among the most abundant human pathogens detected in our samples. Previous research has shown a substantial variety of astroviruses in wastewater samples (Tao et al., 2022; Yang et al., 2021). We therefore aimed to quantify the temporal distribution of different astrovirus subtypes within the investigated timeframe, and to determine the changes in the mutation patterns. As a starting point, we used the taxonomy IDs within the *Astroviridae* family detected by the kaiju metagenomics pipeline (Fig. 2A). For every taxonomy, we selected those sequences (i.e. accession numbers) that matched the collected sequencing data best, as determined by the number of mapped sequencing reads. These sequences were then grouped manually according to their phylogenetic tree (Figure S2A), and one or two sequences per group were selected. Subsequently, point mutations/InDels in the the sequences were then corrected based on the pooled sequencing data to yield the final set, which was again displayed as a phylogenetic tree (Fig. 2B).

**Figure 2.**
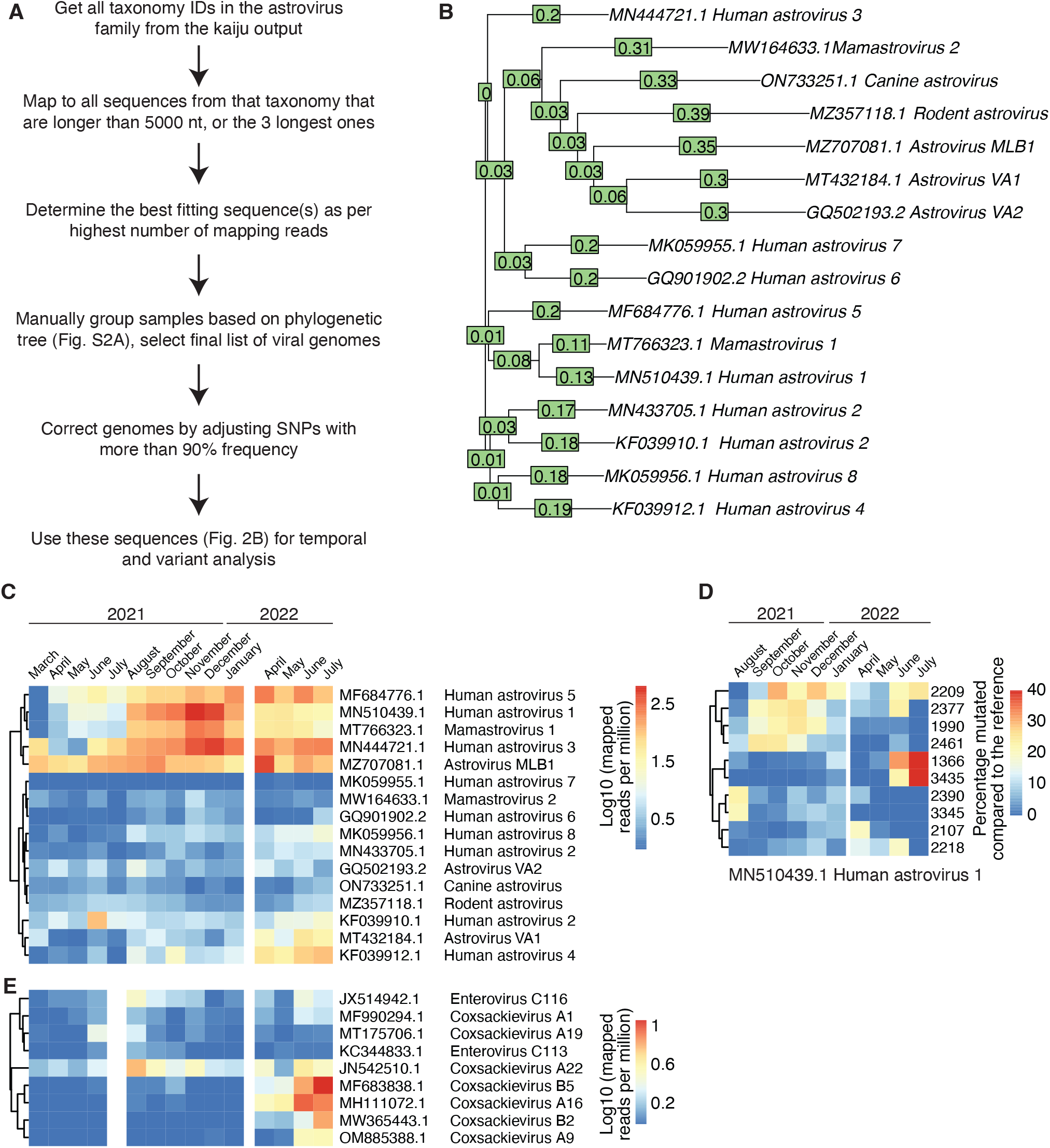
Temporal subspecies/variant dynamics of astroviruses and enteroviruses. **A**, analysis scheme for human astroviruses. **B**, phylogenetic tree of the astroviruses analyzed in detail. **C**, heatmap depicting abundance of the astroviruses over time. Reads are shown as log10 transformed mapped per million, aggreated per month as indicated in the bottom. **D**, for selected position in the human astrovirus 1 genome (based on accession number MN510439.1), the frequency of a mutated residue is shown as a heatmap. **E**, as for C, but for enteroviruses.

We quantified the number of sequences found for the respective accession group per month (Fig. 2C). This analysis showed that the different strains peaked at distinct periods within the investigated time frame. Whereas MLB1 was present throughout the analyzed period, Astrovirus 1 peaked around November/December 2021. Astroviruses VA1 and 4 on the other side emerged only towards the end of the investigation, starting in April 2022.

Since we could obtain full genome coverages over many months for many astroviruses, we were able to quantify mutations per month. As an example, we found variabilities at around 100 positions throughout the Astrovirus 1 (MN510439.1) genome (Fig. S2B). Clustering by variability showed again distinct temporal profiles. A subset of mutated positions is shown in Figure 4D, with e.g. a group of mutations disappearing in fall 2021 (positions 3345, 3390), being only present in fall 2021 (positions 1990, 2461), or emerging in June/July 2022 (positions 1366, 3435).

In addition to profiling known astroviruses, we set out to discover novel ones as well. To do so, all RNA sequencing samples were individually assembled into transcripts using SPAdes (Prjibelski et al., 2020). Next, we searched for assembled transcripts that showed significant sequence similarity to members of the *Astroviridae* family. In total, we found five that could potentially represent novel viruses. Two of them were outside the genera of *Avastrovirus* and *Mamastrovirus*, rather pointing to non-vertebrate hosts (Fig. S2C, upper part). For the three other ones, partial genomes were identified that were grouped within clusters of astroviruses infecting mammals, indicating that some of them could represent novel human pathogens.

### Temporal dynamics of enteroviruses

Next, we analyzed the highly diverse genus of enteroviruses in the sequencing data. This genus contains rhinoviruses, coxsackieviruses, echoviruses, and polioviruses, which can cause a wide range of symptoms, including respiratory illness, meningitis, rash (“hand, foot, and mouth disease”), or paralysis (Harvala et al., 2018). The most important route of transmission is likely via fomites. We followed the same analysis path as for the astroviruses, however we did not apply the genome correction step due to overall low coverage. In general, we observed the expected seasonality (Keeren et al., 2021) with highest signals in northern hemisphere summer months (Fig. 4E). Signal levels were overall lower in the summer of 2021, which could be due hygiene measurements during the SARS-CoV-2 pandemic. Relative abundances were nevertheless distinct, with e.g. subtypes A19 and C113 being mainly present in 2021, and A9, A16, B2 and B5 mainly in 2022.

### Enrichment of specific virus sequences from wastewater

In the previous sections, we focused on viruses that could be readily detected in the total RNA sequencing. However, the signal from many clinically relevant, particularly respiratory viruses was very low or absent. We therefore resorted to a commercial enrichment system (xHYB adventitious agent panel) that can be used to enrich 280 genomes/genome segments from 132 different viruses (see Fig. S3A for a list of viruses). The sequencing libraries generated from total RNA were merged into three pools, as indicated in Table S1. The three pools were then subjected to enrichment, re-amplification, and sequencing. Normalized read counts for all viruses are shown in Fig. S3A, and for a subset with minimal thresholds in Fig. 3. Next to e.g. astroviruses detailed before, this analysis now also captures respiratory viruses, for which the amount of detectable material in the wastewater is low or absent before enrichment (Fig. S3B). This includes respiratory syncytial virus (RSV), influenza, or the common cold coronaviruses NL63, 229E, HKU1, and OC43 (Fig. 3A). We used data from the German Clinical Virology Network (Adams, 2022) in order to relate these observations to clinical diagnostics. Test positivity is shown for a range of respiratory and gastrointestinal viruses (Fig. 3B). A side-by-side comparison for incidence waves of RSV (October/November 2021), Rotavirus A (April-June 2022) or HKU1 (April to June 2022) and NL63 (April 2021 and April 2022), shows the good correlation of data from wastewater and individual testing (Fig. 3C).

**Figure 3.**
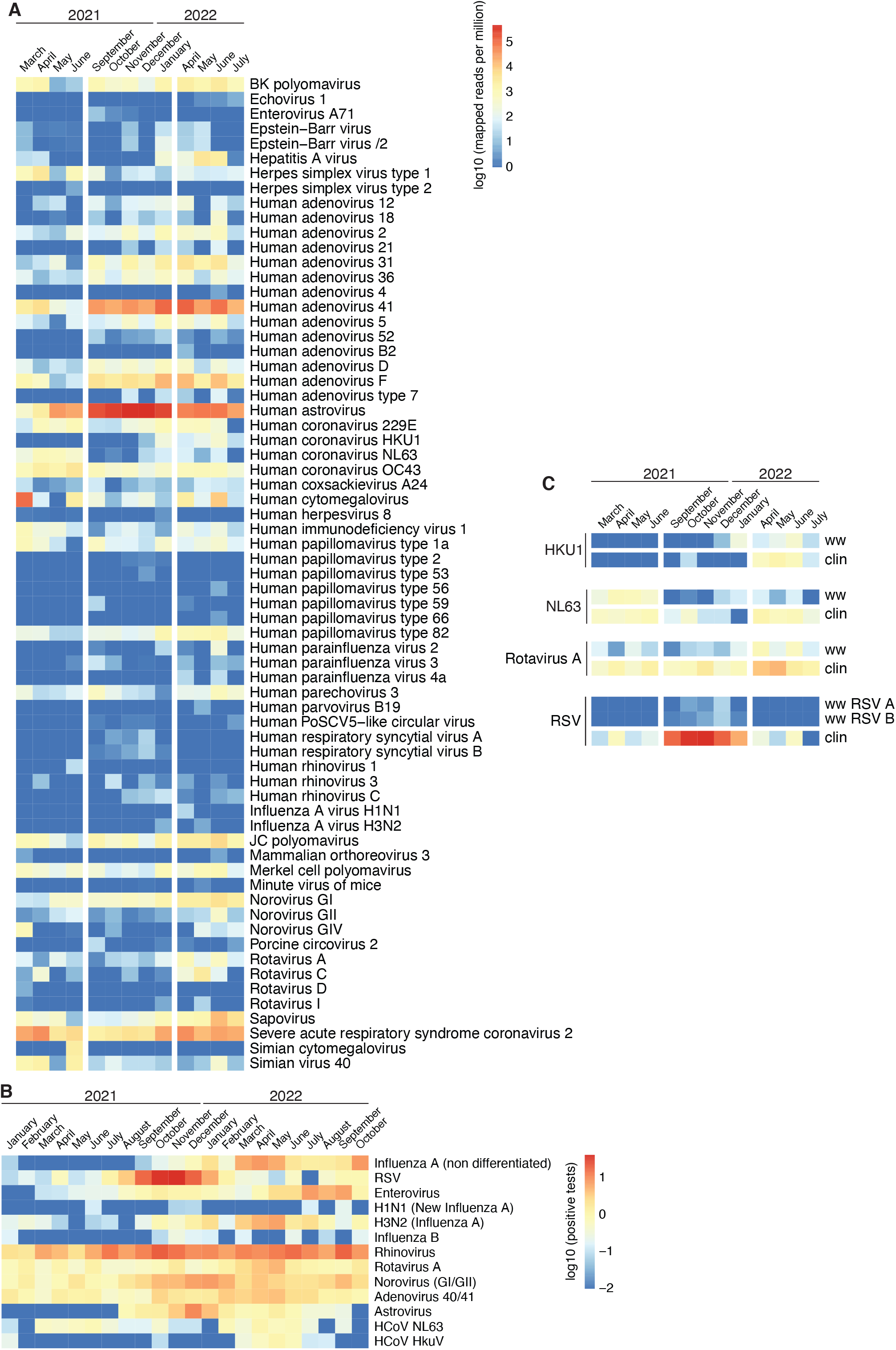
The xHYB enrichment panels captures clinically relevant virus sequences vwith high temporal overlap compared to clinical testing. **A**, heatmap depicting abundance of selected viruses from the enrichment. Reads are shown as log10 transformed mapped per million, aggreated per month as indicated in the bottom, and summed over all parts for segmented genomes. **B**, for selected viruses from the German clinical virology network, shown are log10 transformed percentages of positive tests per month. **C**, direct comparison for four viruses, with for each virus the log10 transformed number of reads mapped per million from wastewater on top (labelled ww), and log10 transformed clinical test positivity (bottom, labelled clin).

The xHYB system enriches on the level of dsDNA, i.e. after the fragmentation steps that are part of the sequencing library preparation protocol. We therefore also tested a procedure that enriches on the level of RNA, starting from the total wastewater RNA. For that purpose, we applied the ElementZero system separately for SARS-CoV-2 and tomato brown mosaic virus, and analyzed the enrichment both using RT-qPCR and high-throughput sequencing. The RTqPCR showed strong depletion of PMMV RNA in all enrichment reactions (about 1000-fold loss, i.e. 10 PCR cycles), and about two- to four-fold loss of the RNA in question, indicating a very strong enrichment (Fig. S3C, upper part). This was corroborated in the sequencing data (Fig. S3C, lower part). Interestingly, when investigating the coverage profiles, we found that SARSCoV-2 showed coverage “islands” in the region covered by hybridization probes, whereas for TBRV the coverage was as equally distributed as for the total RNA, i.e. the input of the ElementZero enrichment (Fig. S3D). An explanation could be that the SARS-CoV-2 RNA is already fragmented in the wastewater, since barely any signal outside of the probe regions was recovered. In contrast, the TBRV RNA would be intact, as the recovery was independent of the distance from the probes.

### Detection of novel viruses from wastewater

Following characterization of known viruses, we set out to search for novel viruses that contain known amino acid sequence motifs, such as the one for the RNA-dependent RNA polymerase. We identified in total 417,972 contigs with sequence similarity to a comprehensive set of known DNA and RNA virus sequences (see Methods for details). Viruses from 49 known und five unclassified (uc) virus orders were discovered. The identified viral sequences were dominated, in terms of number of non-redundant contigs (Fig. 4A, E) and abundance (Fig. 4B, F), by members of the orders *Caudovirales* (DNA viruses), *Norzivirales and Levivirales* (RNA viruses), which infect microbes and likely represent bacteriophages. Many of the contigs were short, with 49,329 (11.8%) being longer than 1000 nt (Fig. 4C, G). Due to the considerable degree of sequence fragmentation we cannot exclude that the actual number of viruses is lower than the number of reported viral contigs as a virus might contribute with several sequence fragments. The majority of 277,092 of the viral contigs (66.3%) showed a protein sequence identity to the closest known virus of less than 90% (Fig. 4D, H), indicating that these sequences originated from novel viruses.

**Figure 4.**
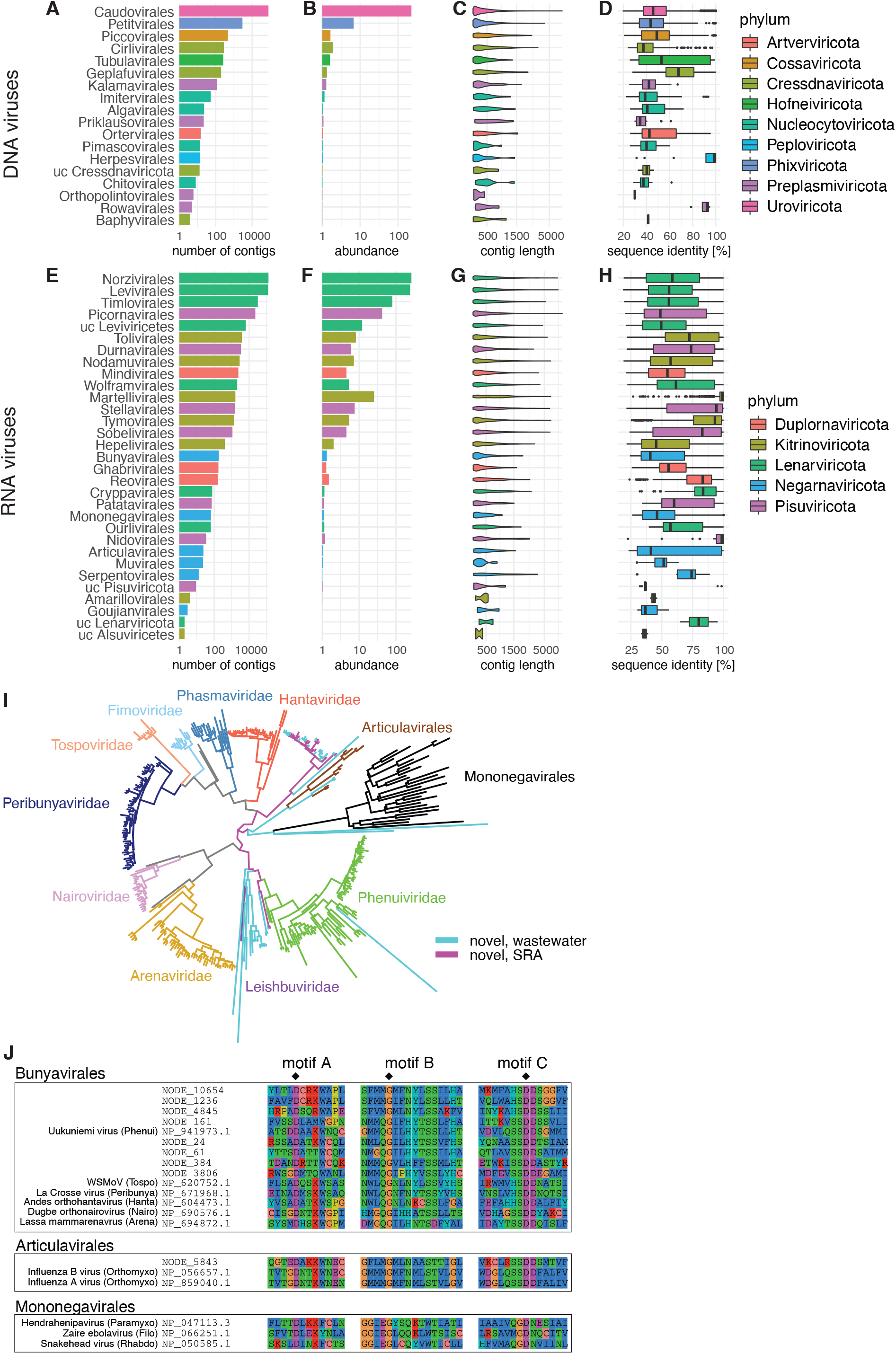
Novel viral sequences discovered from wastewater. **A, E**, Number of non-redundant novel DNA viral (top row) and RNA viral (bottom row) contigs identified in all waste water samples together, aggregated by order and colored by phylum. **B, F**, Viral abundances. **C, G**, Lengths of the contigs within the order labelled on the left. **D, H**, Protein sequence identities to the closest known reference virus for the orders labelled on the left. **I**, L protein-based Bayesian phylogeny of known and novel bunya- and articulaviruses. Bunyavirus families, the Articula-virales order and the outgroup Mononegavirales order (black) are discriminated by branch color. Viruses discovered from wastewater and the SRA screen are in cyan and purple, respectively. **J**, Multiple sequence alignment of the region around most conserved motifs A, B and C of the viral RdRp for a selected set of novel and reference viruses. Two catalytic residues of motifs A and C and one residue of motif B involved in nucleotide selection are marked by diamond symbols.

Among the viral sequences, 107 were novel, non-redundant sequences that were classified as *Bunyavirales* or its sister order *Articulavirales*. These included sequences that prototype distant lineages within the *Bunyavirales* in the L protein-based phylogeny (Fig. 4I), suggesting that some of the newly discovered viruses form novel virus families. The majority of the contigs were identified via sequence homology to L proteins of known bunya- and articulaviruses while others gave hits against nucleocapsid proteins or glycoproteins of certain bunyavirus families. To provide additional and independent evidence that the discovered viruses are genuine, we retrieved 16 bunyavirus-like sequences that shared 80% or more protein sequence identity to the viruses identified from wastewater and which we discovered in a large screen of public transcriptome projects from the Sequence Read Archive (SRA) (Fig. 4L). The fact that we rediscovered some of the viral sequences from the wastewater analysis in the SRA analysis provides independent support for the authenticity of the described novel viruses. An amino acid sequence alignment of motifs A, B and C of the RNA dependent RNA polymerase shows conserved regions among the subfamilies to which the novel viruses belong (Fig. 4J). Finally, we probed the amount of five of the most abundant novel virus contigs across the entire sampling time course. Three of the five were detected in only one sample each, one over the entire time course, and the fifth was restricted to the period from April to May 2022 (Fig. S4A).

## Discussion

In this study, we present a deep longitudinal profiling of the wastewater virome from a treatment plant in Berlin, Germany. Overall, we analyzed samples covering a period of 17 months, from March 2021 to July 2022. Of note, at the beginning of the time series, there were still substantial measures for mitigation of viral spread in place due to the SARS-CoV-2 pandemic, which very likely also affected a range of pathogens. From May 2021 onwards, there was a mask mandate in place, however no closures or shutdown measures. The data therefore may differ from a situation without such measures, as before the pandemic.

Nevertheless, we could track a wide variety of microbes, including a broad range of human viruses. Previous wastewater metagenomics studies (Adriaenssens et al., 2018; Bibby and Peccia, 2013; Cantalupo et al., 2011; Fernandez-Cassi et al., 2018; Guajardo-Leiva et al., 2020; Martinez-Puchol et al., 2021; Perez-Cataluna et al., 2021; Rothman et al., 2020; Rothman et al., 2021) have assessed the scope of taxa to be found in this sample type. These findings, such as the dominance of bacterial sequences as well as the high abundance of plant viruses were recapitulated in our data. In extension of this work, we present several findings. First, the relatively broad sampling period allowed the detection of non-human seasonal viruses such as those in seasonal food (grapevines, watermelons) as well as in mosquitos, which are in Berlin present only during Northern hemisphere summer (Fig. 1C). Such findings provide some degree of insurance for the seasonality in our data set, but also hint to the breadth of the information that wastewater can contain, to allow monitoring of entire ecosystems, and not merely specific pathogens. Second, our detailed and temporal investigations of astrovirus and enterovirus subspecies, as well as the temporal variant dynamics exemplified in mamastrovirus 1 (Fig. 2) shows the depth of information which can be recovered by wastewater monitoring. Third, although several clinically relevant respiratory viruses are – not surprisingly, in contrast to gastroenteritis viruses – much less abundant, they can be recovered in high-throughput sequencing using targeted enrichment approaches (Fig. 3). Comparison with clinical testing obtained from individual patients shows high agreement (Fig. 3C), underlining that wastewater can, with appropriate methodology, possibly be used to track every circulating human pathogen. An interesting observation was that enrichment on the RNA level recovered only highly fragmented genome coverage from the enveloped virus SARS-CoV-2. This finding indicates that at least this RNA, and maybe RNA from enveloped viruses in general, is present not as part of viral particles but rather bound to protein proteins or other matter (Fig. S3). Along with observations such as the higher abundance of mpox and influenza DNA/RNA in the particle fractions (Wolfe et al., 2022a; Wolfe et al., 2022b), this underscores the need for systematic investigations into which processing methods are suitable for the detection of a specific microbe. And fourth, our discovery of possibly tens of thousands novel viruses shows that wastewater can also fill major gaps in our knowledge of the planetary virome and thus can inform future assessment of zoonotic potentials.

Our study has several limitations. We have only investigated one wastewater treatment plant at one location, albeit longitudinally. Together with increasing deep wastewater metagenomic data, global comparisons will become possible. Furthermore, our processing method may capture a wide array of microbes, however it certainly misses others. For example, despite relatively high local incidences, we have not been able to detect mpox genomes in our samples in contrast to other sites (Wannigama et al., 2023; Wolfe et al., 2022b). Also, temperature differences (in Berlin, wastewater is in the range between 15 °C in winter and 25 °C) as well as the flow time from the households to the treatment plant (in the range of one to several hours) can introduce variability that is difficult to track. And finally, although our sampling series is to our knowledge the longest published so far, it still captures seasons only once or twice, and in parts was done during an exceptional period of pandemic mitigation measures. With still decreasing costs for high-throughput sequencing along with refined experimental and computational methodology however, future studies will be able to capture microbial communities at a so far unfathomable level of detail.

## Acknowledgements

We are very grateful to Fredrik Zietzschmann, Katharina Flatau, Regina Gnirss and Uta Böckelmann from the municipal water authority (Berliner Wasserbetriebe) for providing samples and continuous support. Figure 1A was generously created and provided by Thomas Zahn, Peter Pennitz and Martin Witzenrath. We thank Sindy Böttcher and Julian Kreibich for support particularly with enteroviruses, as well as René Kallies, Antonis Chatzinotas, Hans-Christoph Selinka and Martin Meixner for support and helpful discussions. EW, CL, GTA, SS, and ML are supported by the Project “Virological and immunological determinants of COVID-19 pathogenesis – lessons to get prepared for future pandemics (KA1-Co-02 ‘COVIPA’)”, a grant from the Helmholtz Association Initiative and Networking Fund. CL acknowledges support by the Deutsche Forschungsgemeinschaft (DFG, German Research Foundation) under Germany’s Excellence Strategy - EXC 2155 - project number 390874280. CL is a member of the European Virus Bioinformatics Center (EVBC).

## Author contributions

EW, AD, CQ and GTA performed experiments. EW, CL, AM, SS and ML analyzed data and prepared figures. EW wrote the manuscript, with contributions from CL. TB, JA and SS supervised parts of the projects, ML supervised the entire project. All authors approved the manuscript.

## Data and code availability

All raw sequencing data will be made available via NCBI GEO, and code on github.com, shortly. Meanwhile, please contact for access.

