## Supplementary Figures for "Comprehensive profiling of wastewater viromes by genomic sequencing"

#### Figure S1, part 1

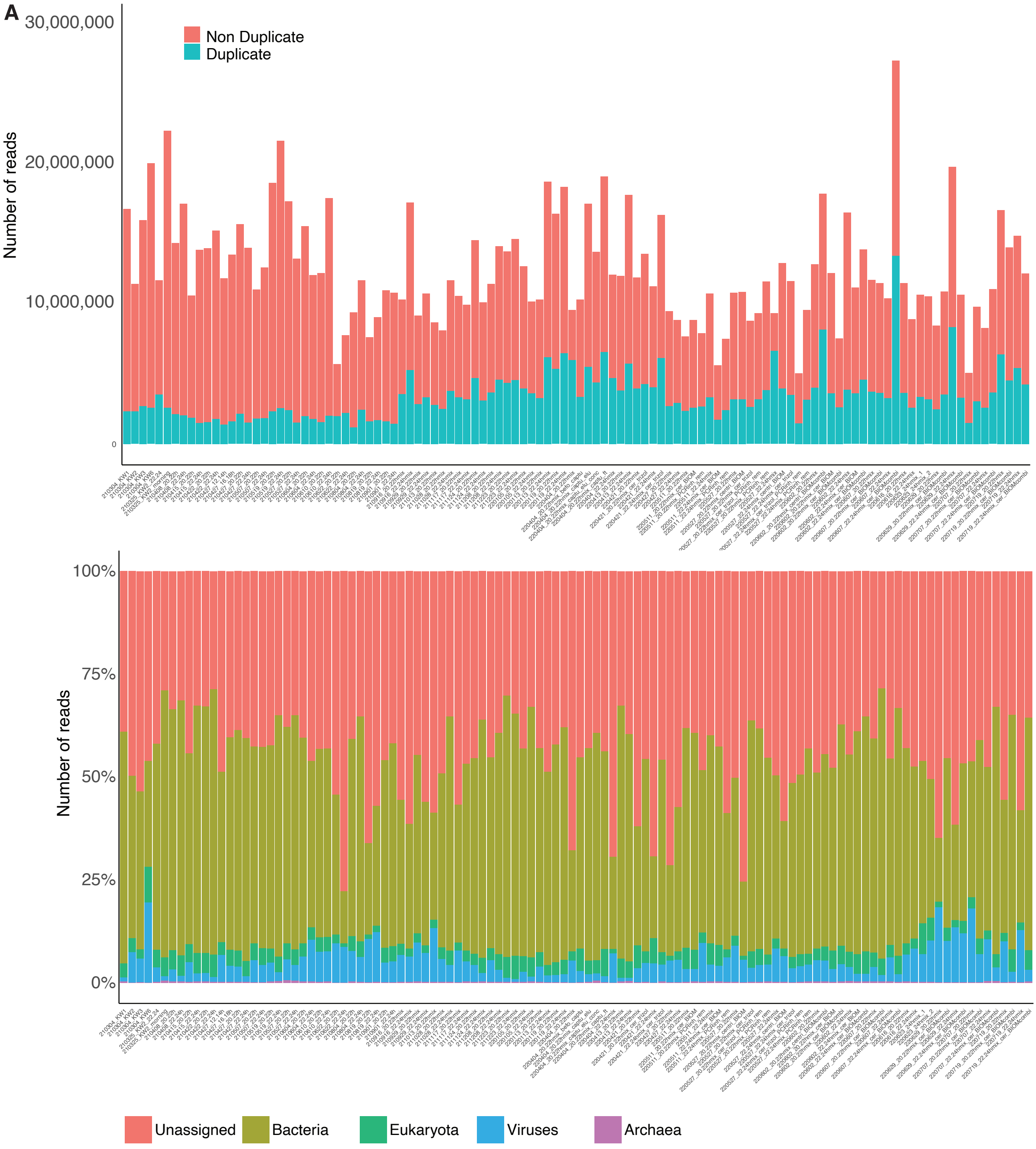


**Figure S1. A,** number of raw reads per sample, with non duplicated reads in red, and duplicates reads in petrol. **B,** percentages of kaiju assignements to bacteria/eukaryota/viruses/archaea, as well as the percentage of reads that could not be assigned to a taxonomy. **C,** Princial component analysis (PCA) of all samples based on the 16082 taxonomies after initial filtering, with outlier samples labeled, and the top three contributors to PC1 and PC2, respectively, labelled with red arrows as well as taxonomy ID and name. **D,** barplot with the number of reads assigned to the outlying annotations (Methods). **E,** heatmap depicting abundance of the top 20 families for viruses, eukaryotes and bacteria, and 14 families for archeae. Reads are shown as log10 transformed mapped per million, aggreated per month as indicated in the bottom. The bottom panel shows 0-1 normalized signal for four specific viruses. **F,** as in Fig. 1A but for families.

Figure S1, part 2

**C**

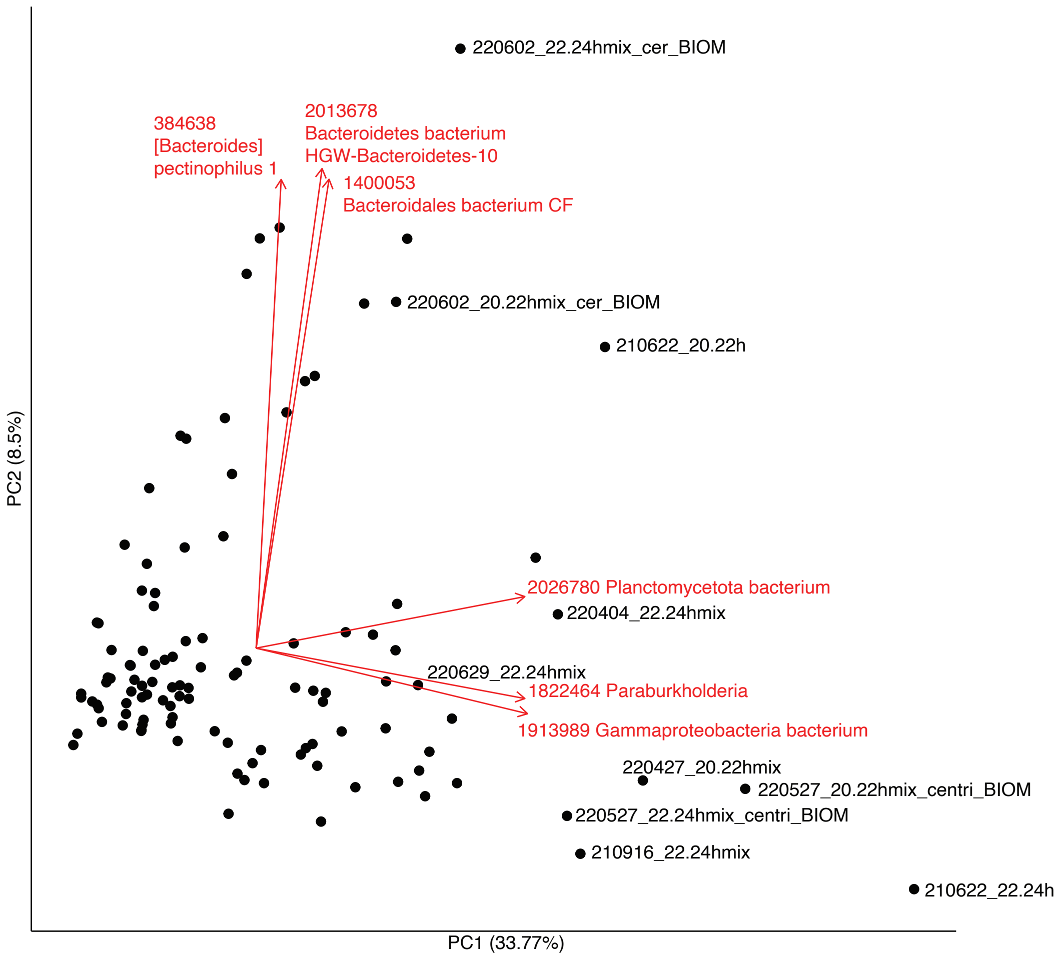


**D**

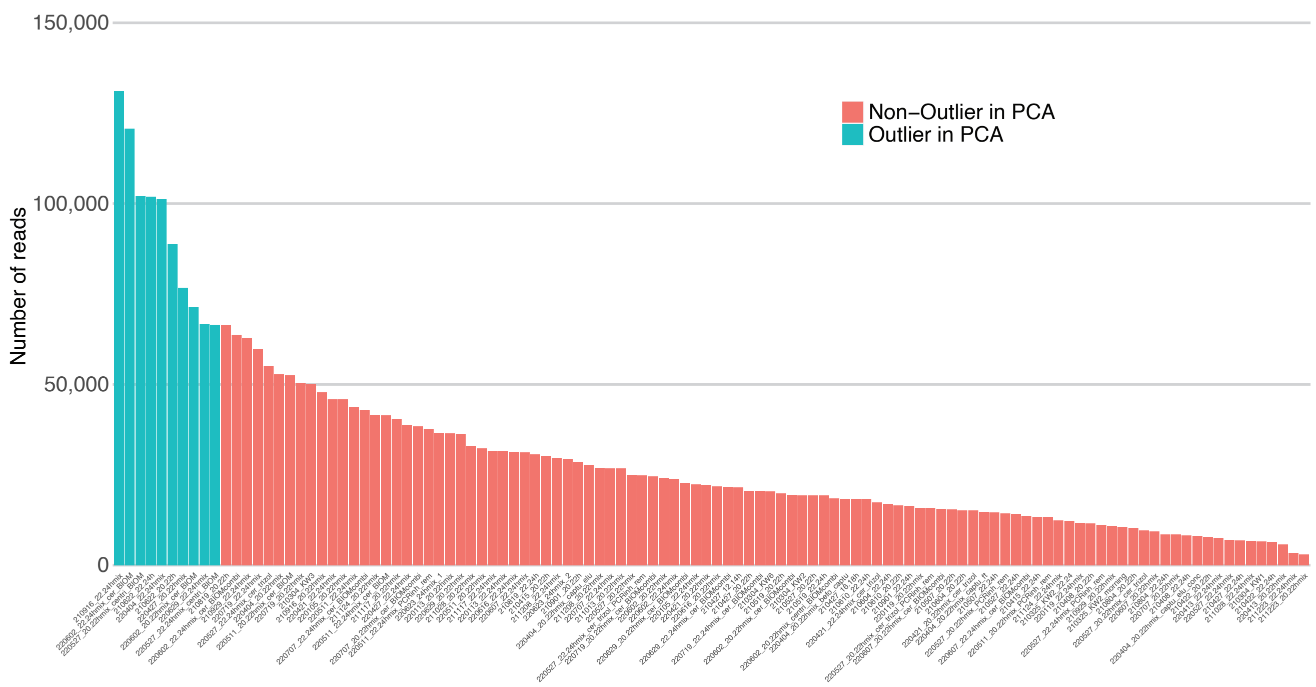

Figure S1, part 3

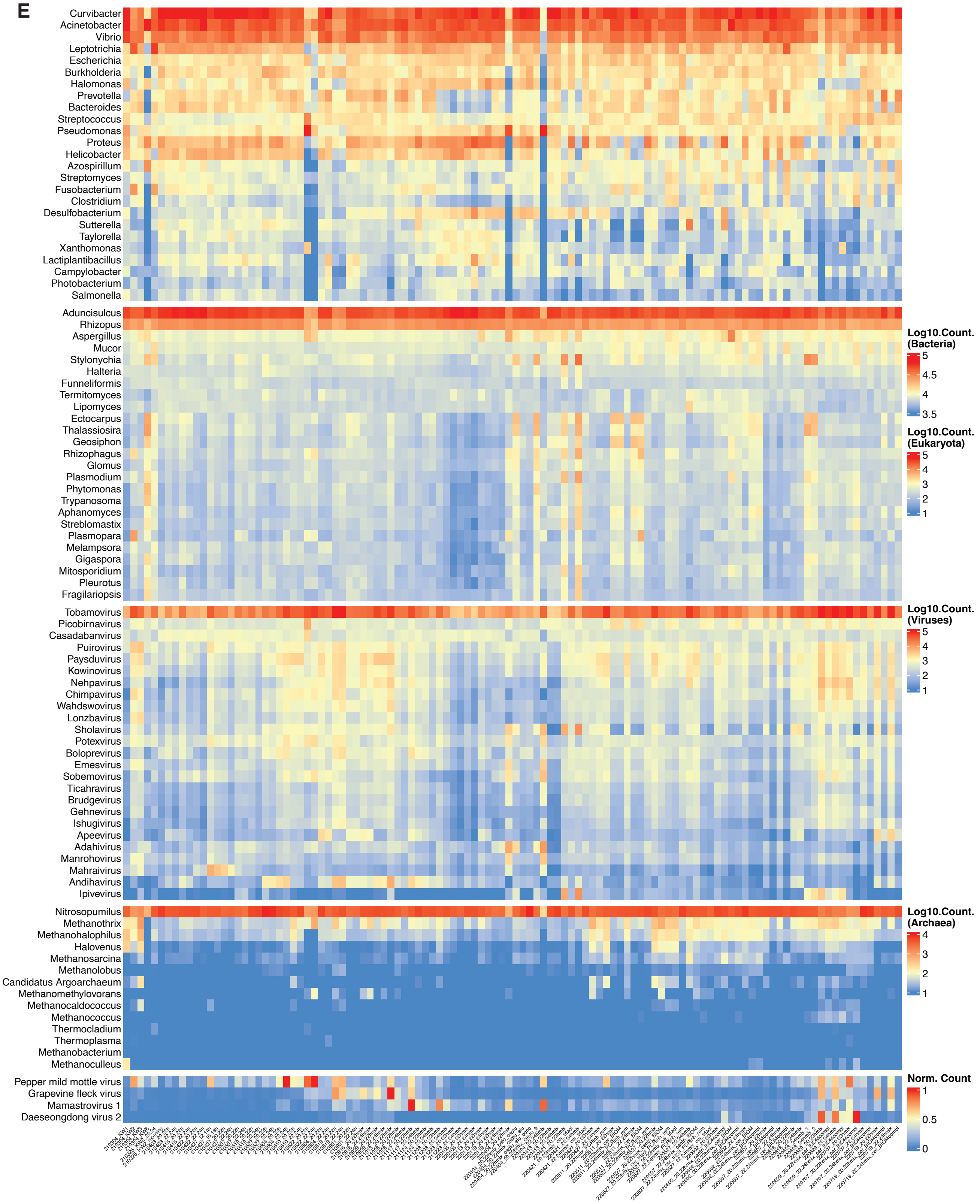

Figure S1, part 4

**F**

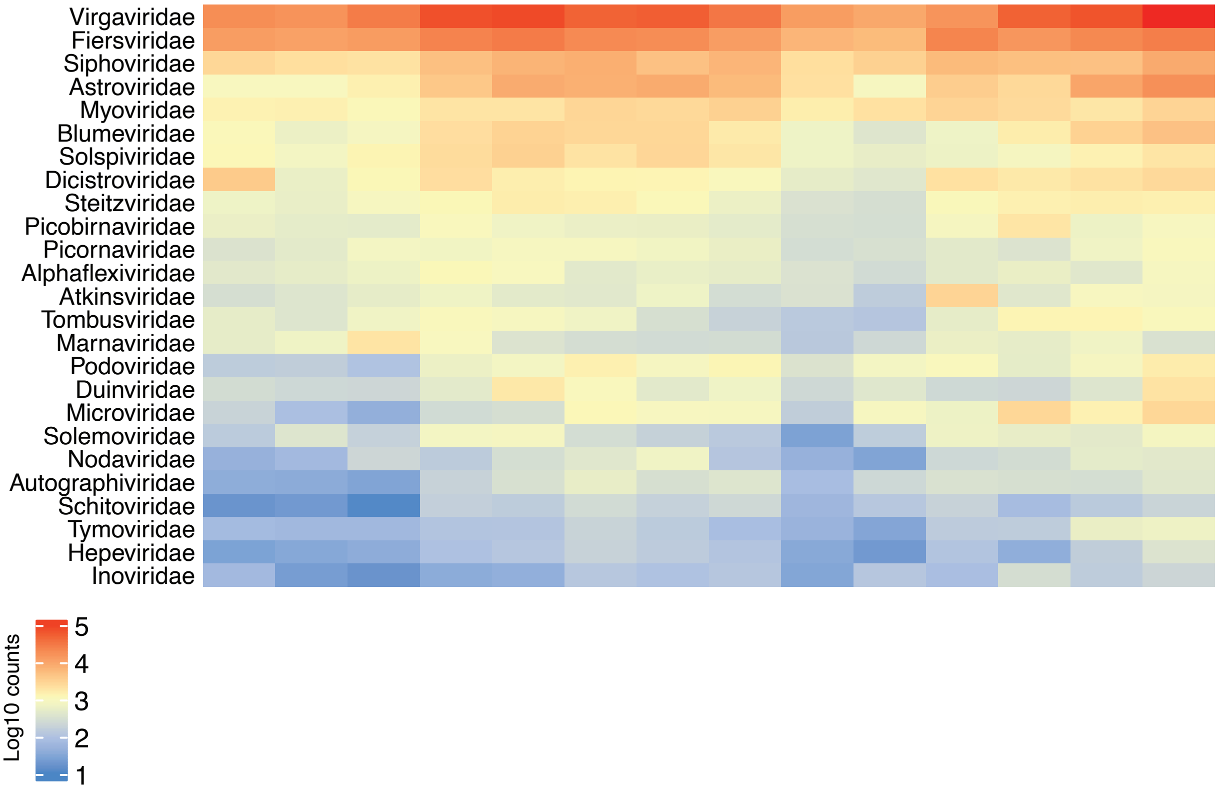

Figure S2, part 1

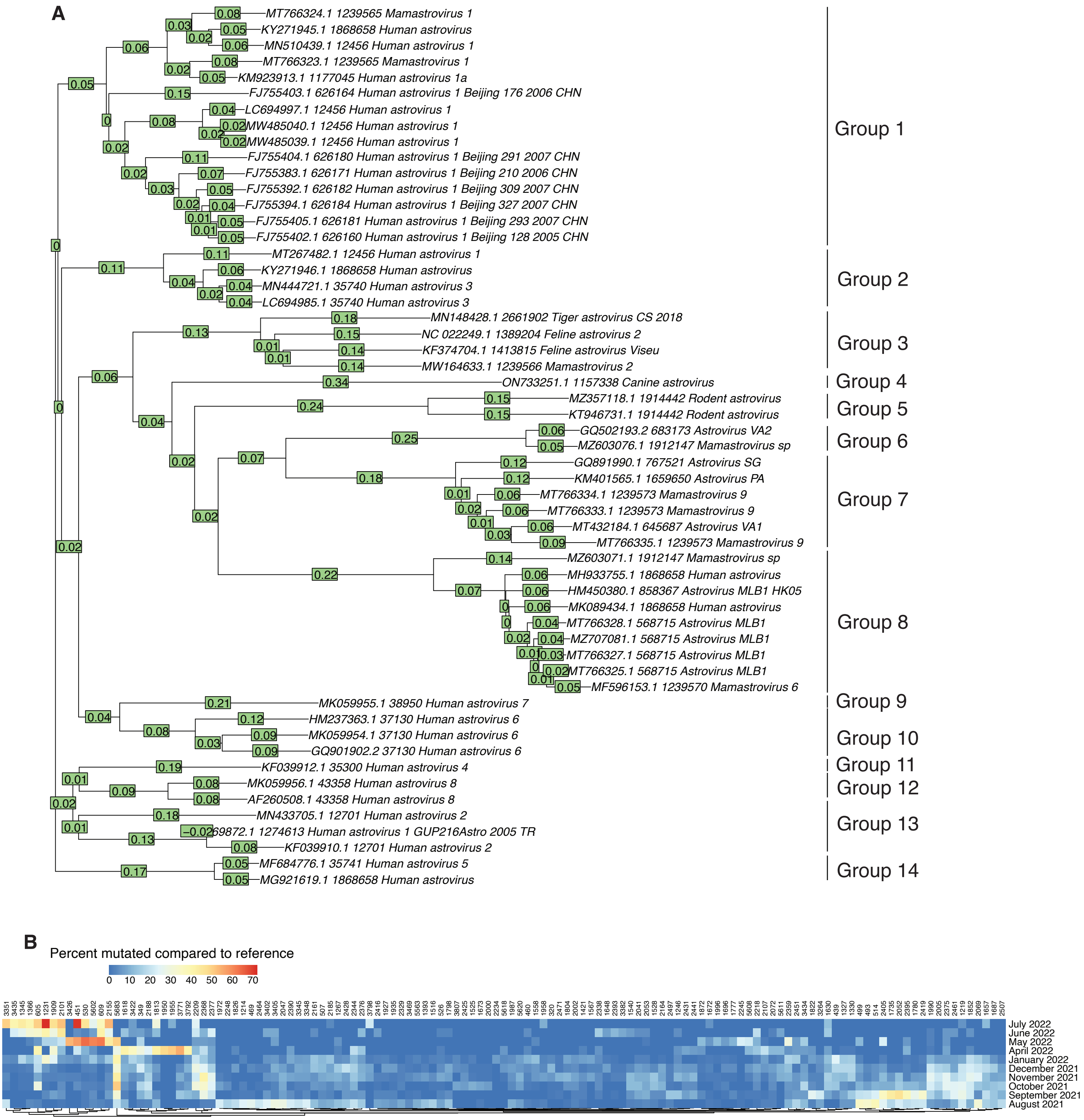


**Figure S2. A,** phylogenetic trees of the sequences from the accession numbers that showed closest proximity to the sequencing data in this study. Manually assigned groups, from which 1 or 2 sequences were selected for further analysis, are shown on the right. **B,** for all positions in the human astrovirus 1 genome (based on accession number MN510439.1) with minimally 5% mutation frequency in at least one sample, the frequency of a mutated residue is shown as a heatmap. **C,** Shown is an RdRp-based Maximum Likelihood phylogeny of reference astroviruses and five novel astro- and astro-like viruses (names in red) with contig lengths of at least 1000 nt and protein sequence identity to the closest reference of less than 90%. Bastroviruses are used as an outgroup, Avastroviruses are shaded in blue, mamstroviruses in yellow.

Figure S2, part 2

**C**

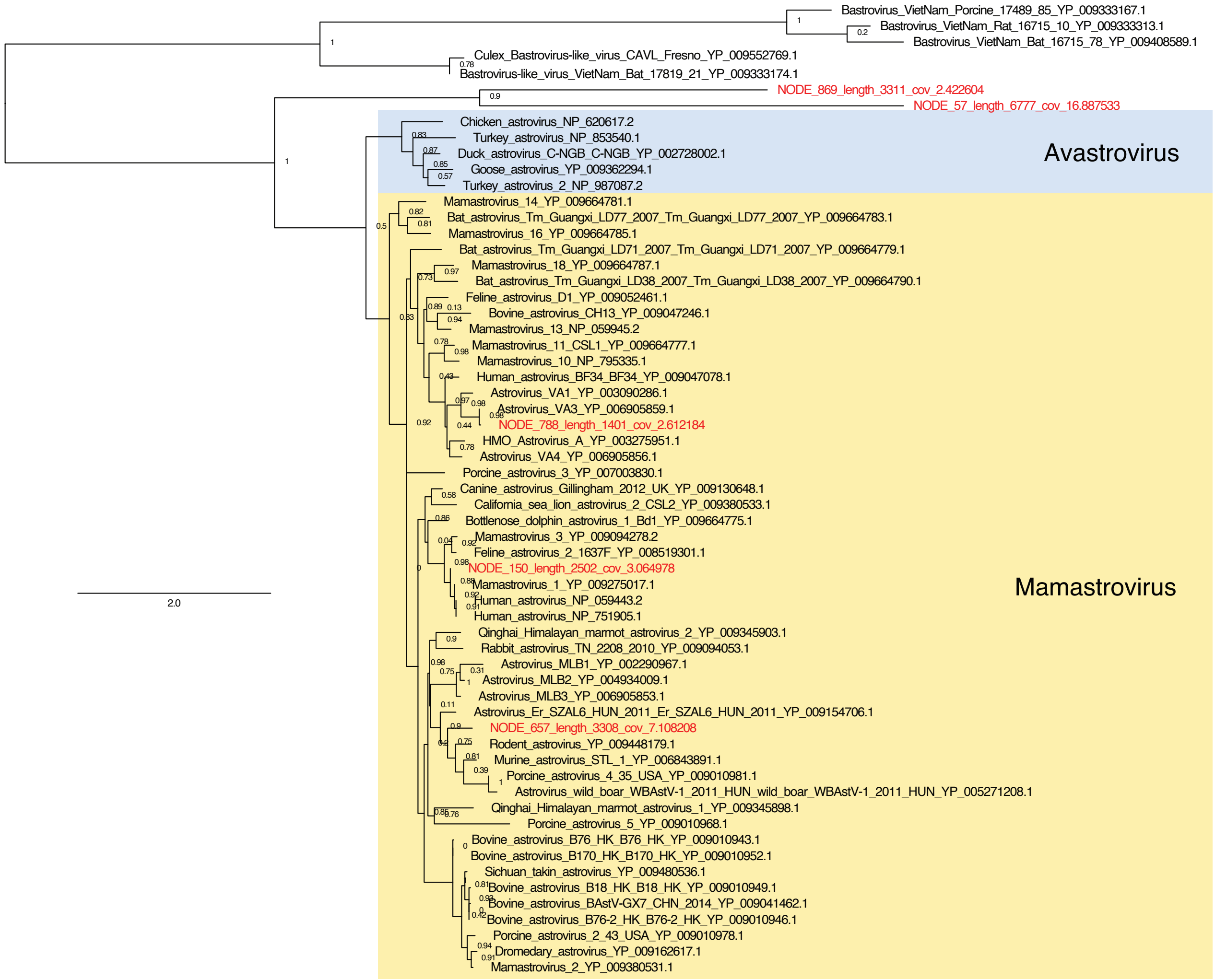

### Figure S3, part 1

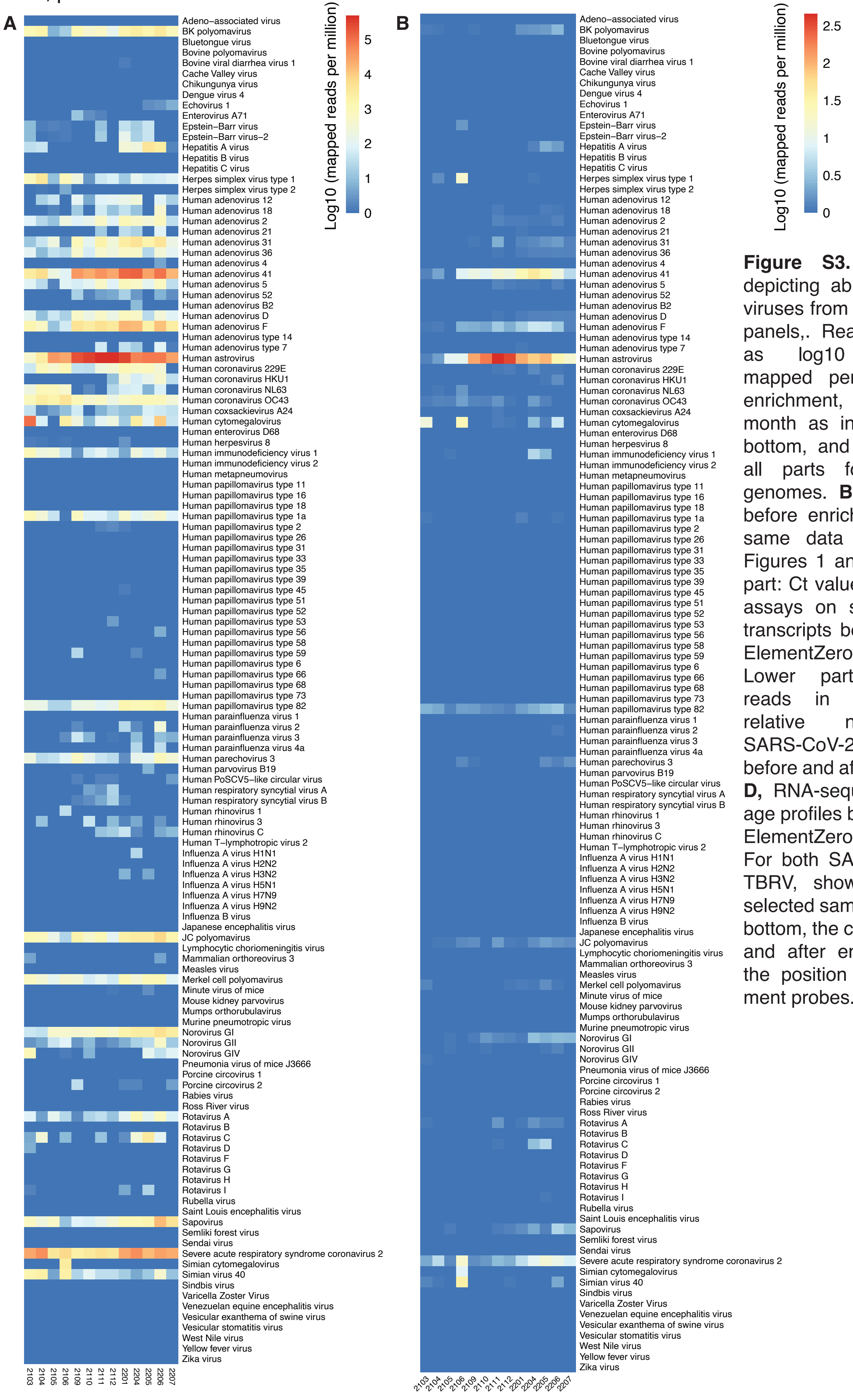


**Figure S3. A,** heatmap depicting abundance of all viruses from the enrichment panels,. Reads are shown as log10 transformed mapped per million after enrichment, aggreated per month as indicated in the bottom, and summed over all parts for segmented genomes. **B,** as in A, but before enrichment (i.e. the same data as shown in Figures 1 and 2). **C,** upper part: Ct values of RT-qPCR assays on selected target transcripts before and after ElementZero enrichment. Lower part, sequencing reads in absolute and relative numbers for SARS-CoV-2 and TBRV before and after enrichment. **D,** RNA-sequencing coverage profiles before and after ElementZero enrichment. For both SARS-CoV-2 and TBRV, shown are, for a selected sample, from top to bottom, the coverage before and after enrichment, and the position of the enrichment probes.

#### Figure S3, part 2

##### C RT-qPCR, Ct values

| | | | PMMV | PMMV | E | E | N1 | N1 | N2 | N2 | IMP-Orf1b | IMP-Orf1b | TBRV | TBRV |
|---|---|---|---|---|---|---|---|---|---|---|---|---|---|---|
| ElementZero eluate | input | WW0119_20-22h mix | 25,3 | 25,3 | 33,1 | 33,2 | 33,4 | 34,7 | 33,2 | 35,3 | 34,3 | 33,8 | 20,9 | 20,8 |
| | | WW0119 22-24h mix | 24,4 | 24,4 | 33,4 | 33,2 | 32,8 | 33,2 | 33,8 | 33,6 | 33,4 | 34,3 | 23,8 | 24,0 |
| | | WW0204 22-24h mix | 26,1 | 26,2 | 34,4 | nd | nd | 34,5 | nd | 35,2 | 34,6 | 34,8 | 22,9 | 22,7 |
| | SARS-CoV-2 | WW0119_20-22h mix | nd | 35,6 | nd | 35,8 | nd | nd | nd | 36,5 | nd | nd | nd | nd |
| | | WW0119 22-24h mix | 35,3 | 35,4 | 35,3 | nd | nd | 36,6 | nd | nd | 35,5 | nd | 36,3 | nd |
| | | WW0204 22-24h mix | 35,2 | 36,3 | nd | 37,0 | nd | 37,1 | nd | nd | nd | nd | nd | nd |
| | TBRV | WW0119 20-22h mix | 35,3 | 35,7 | nd | nd | nd | nd | nd | nd | nd | nd | 23,7 | 23,7 |
| | | WW0119 22-24h mix | 34,2 | 36,3 | nd | nd | nd | nd | nd | nd | nd | nd | 22,3 | 22,8 |
| | | WW0204 22-24h mi | 34,4 | 36,9 | nd | nd | nd | nd | nd | nd | nd | nd | 24,9 | 25,3 |

nd = not detected

sequencing

| | | | | aligning reads | | aligning reads, per million | |
|---|---|---|---|---|---|---|---|
| | | | total reads | SARS-CoV-2 | TBRV | SARS-CoV-2 | TBRV |
| ElementZero eluate | input | WW0119_20-22 | 16634889 | 12 | 66331 | 1 | 3987 |
| | | WW0119 22-24 | 18592825 | 9 | 102505 | 0 | 5513 |
| | | WW0204 22-0 | 6265867 | 9 | 38831 | 1 | 6197 |
| | SARS-CoV-2 | WW0119_20-22h mix | 6621905 | 2927 | 93 | 442 | 14 |
| | | WW0119 22-24h mix | 8730960 | 3020 | 331 | 346 | 38 |
| | | WW0204 22-24h mix | 7443317 | 2729 | 27 | 367 | 4 |
| | TBRV | WW0119 20-22h mix | 5731701 | 0 | 1653218 | 0 | 288434 |
| | | WW0119 22-24h mix | 10717491 | 1 | 743102 | 0 | 69335 |
| | | WW0204 22-24h mix | 7734757 | 2 | 483384 | 0 | 62495 |

### D

Sample 220119 22-24hmix

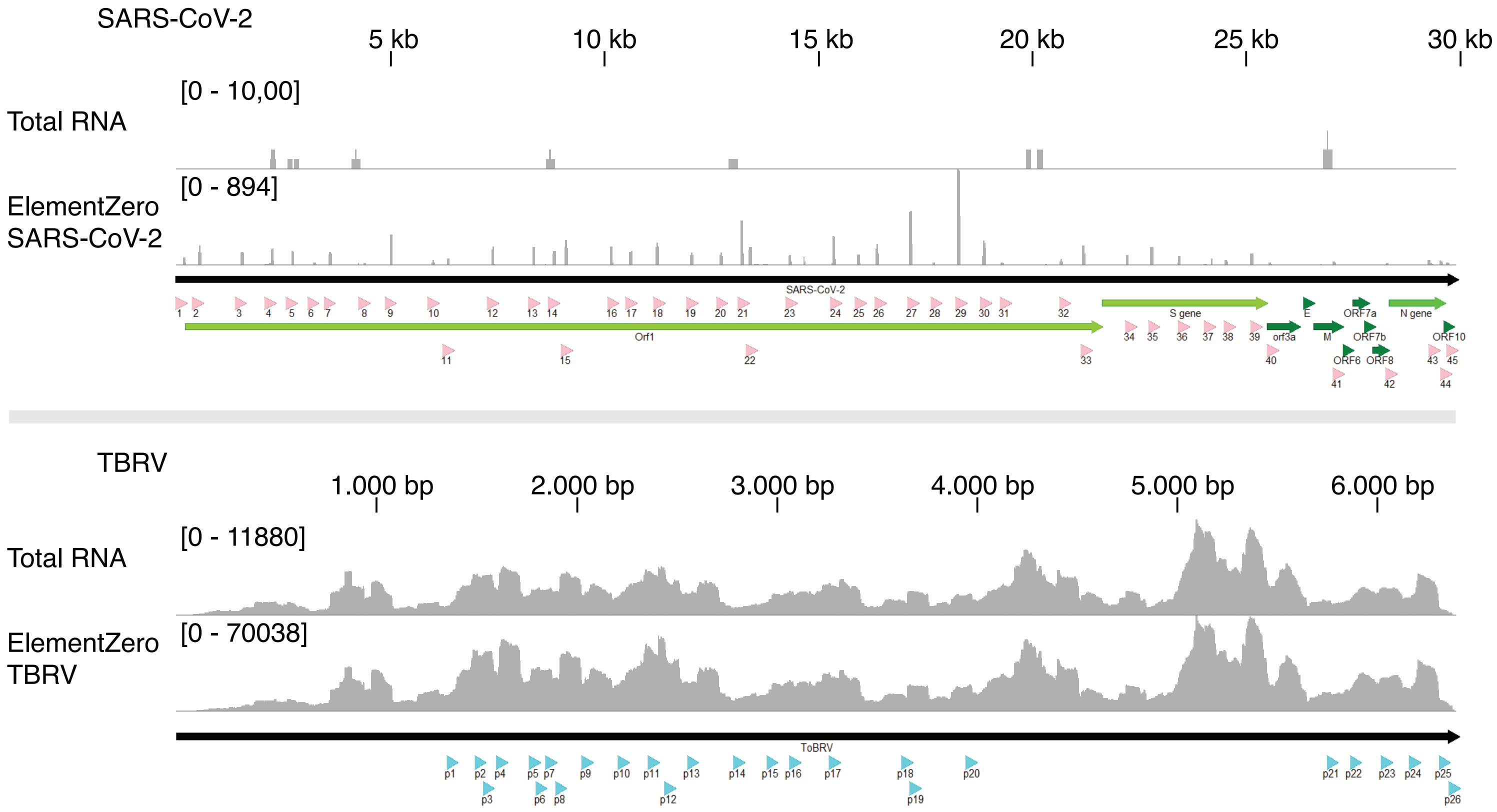

Figure S4

**A**

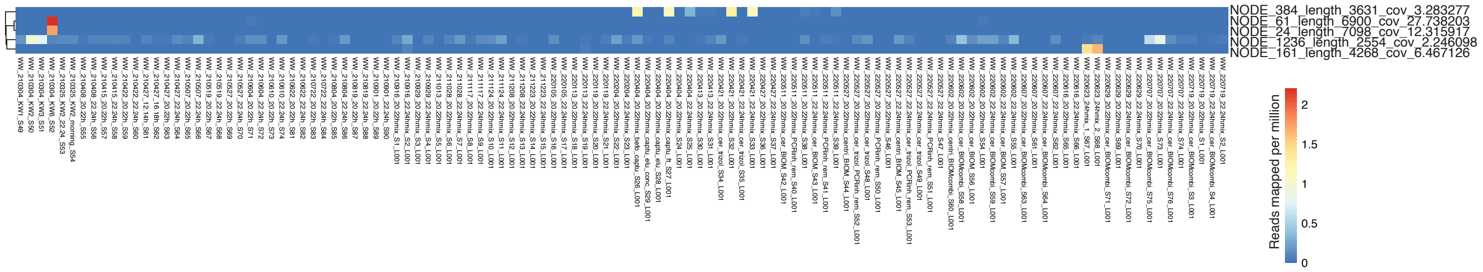
NODE_384_length_3631_cov_3.283277
NODE_61_length_6900_cov_27.738203
NODE_24_length_7098_cov_12.315917
NODE_1236_length_2554_cov_2.246098
NODE_161_length_4268_cov_6.467126
Reads mapped per million
2
1.5
1
0.5
0
